# A conserved lamination pattern in the paleocortex and the neocortex revealed by single-cell RNA analyses

**DOI:** 10.64898/2026.07.02.732788

**Authors:** Makoto Nasu, Shigeyuki Esumi, Tomohiko Wakayama, Kenji Shimamura

## Abstract

The piriform cortex, the largest paleocortical domain and a central olfactory cortex, is classically described as a three-layered structure, with layer II subdivided into semilunar and superficial pyramidal neurons. However, its cellular composition, developmental logic, and evolutionary relationship with the six-layered neocortex remain incompletely defined. Here, we aimed to resolve piriform cortical cell types and laminar organization and compare their molecular programs across mammalian cortical regions and the reptilian cortex. We performed single-nucleus RNA sequencing of the microdissected piriform cortex and integrated neuronal lineage data with published datasets, with spatial validation via the Allen Mouse Brain Atlas. Interspecies analyses identified piriform-dominant glutamatergic populations marked by Unc13c/Lmo3/Rora and an Rorb/Reln/Ntng1-enriched sensory-recipient subtype localized to the superficial layer IIa, consistent with semilunar neurons. Spatial transcriptomic mapping revealed that piriform layer IIb contains neocortical upper-layer-like corticocortical neurons, while piriform layer III segregates into layer V–like (IIIa), layer VI–like (IIIb), and VIb/VII (subplate)-equivalent populations, indicating a neocortex-like laminar framework with an inverted positioning of sensory recipients and corticocortical compartments relative to the neocortex. GABAergic neurons largely conformed to canonical MGE- and CGE-derived lineages but included a piriform-specific Rgs9+Pde7b+ subtype consistent with an LGE-related origin, suggesting region-specific diversification of GABAergic neurons. Disease ontology enrichment linked hippocampus-dominant glutamatergic programs to Alzheimer’s-related genes and piriform-dominant (and pan-cortical) excitatory and inhibitory programs to autism spectrum disorder and intellectual disability, implicating coordinated E/I circuit specializations in area-selective vulnerability. These findings support a revised view of the paleocortex as a divergent specialization of a conserved, multilayered cortical program and provide molecular markers and a comparative framework for studying cortical evolution and region-specific disease susceptibility.

## Introduction

The piriform cortex (Pir), the largest domain in the paleocortex (PCx), lies lateral to the neocortex (NCx) and primarily functions as a central olfactory cortex (Puelles *et al*., 2000; Butler and Molnár, 2002; García-Moreno *et al*., 2008; Medina and Abellán, 2009). Although the Pir is one of the sensory cortices, it exhibits several distinctive features compared to the NCx. In particular, Pir displays a three-layered structure—layer I (molecular layer), layer II (pyramidal layer), and layer III (polymorph layer)—in contrast to the six-layered NCx (Luzzati, 2015; Shepherd and Rowe, 2017; Tosches and Laurent, 2019). Traditionally, layer II of the Pir has been subdivided into a superficial layer IIa and a deeper layer IIb, comprising two morphologically and electrophysiologically distinct neuronal types: semilunar (SL) neurons in layer IIa and superficial pyramidal (Pyr) neurons in layer IIb (Haberly and Price, 1978; Suzuki and Bekkers, 2006). SL neurons preferentially input sensory information from the olfactory bulb (OB) and output to the posteromedial cortical amygdala and lateral entorhinal cortex (Diodato *et al*., 2016; Mazo *et al*., 2017; Nagappan and Franks, 2021; Chen *et al*., 2022b). In contrast, Pyr neurons receive both olfactory bulb afferents and associational/commissural inputs from other cortical areas and project back to the OB, medial prefrontal cortex, and orbitofrontal cortex. These studies provide some insight into Pir-involving cerebral circuitry; however, the Pir has hardly been considered as the responsible site of neurological diseases and major cortical functions.

From an evolutionary and developmental perspective, the PCx has long attracted attention as a key structure for exploring the origin of the six-layered neocortex, which is a defining feature of the mammalian pallium. In contrast to the nuclear organization of the avian pallium and the partially three-layered cortex of reptiles, the simpler laminar architecture of the PCx occupies an intermediate position that may reflect the ancestral developmental programs of cortical patterning (Butler and Molnár, 2002; Medina and Abellán, 2009; Butler *et al*., 2011; Karten, 2013; Nomura *et al*., 2014; Montiel *et al*., 2015; Shepherd and Rowe, 2017; Briscoe and Ragsdale, 2018; Tosches, 2021; Zaremba *et al*., 2025).

Although the Pir expresses a characteristic repertoire of region-specific genes (Medina and Abellán, 2009; Diodato *et al*., 2016; Nasu *et al*., 2020; Hara *et al*., 2025; Zeppilli *et al*., 2025), its cellular composition and developmental logic remain incompletely defined, leaving its evolutionary position within the pallium unresolved. To address this issue, we conducted transcriptomic profiling of Pir from 3-week-old mice. In addition, we conducted spatial and evolutionary comparisons across cortical regions to identify features unique to the piriform cortex and cellular programs shared with other cortical areas.

## Results

### RNA-seq of the paleocortex

To characterize the cellular diversity of the mouse PCx, we performed single-nucleus RNA sequencing (snRNA-seq) on the Pir of 3-week-old mice, a developmental stage at which neuronal differentiation is largely complete and secondary physiological influences are expected to be minimal (Figure 1A). After quality control, we obtained high-quality transcriptomic profiles from 8,822 nuclei (Supplementary Figure 1A–D). Reference-based classification and dimensionality reduction by UMAP identified three major classes: glutamatergic neurons (Glu), GABAergic neurons (GABA), and non-neuronal glial cells (Glia) (Hao *et al*., 2021). Based on a reference dataset derived from the mouse primary motor cortex (Yao *et al*., 2020), our dataset contained 6,131 Glu, 954 GABA, and 1,737 glia (Supplementary Figure S2A,B). Most neurons were clearly separated from Glia, while Glu and GABA clusters were positioned in proximity, with some neuronal lineage cells intermingling with glial populations. The neuron-to-glia ratio was 3.8:1, which is comparable to that of the mouse NCx (Zeisel *et al*., 2015), suggesting that by 3 weeks postnatally, the Pir had reached a developmental stage resembling that of the NCx, where neurogenesis was complete and gliogenesis had begun.

**Figure 1.**
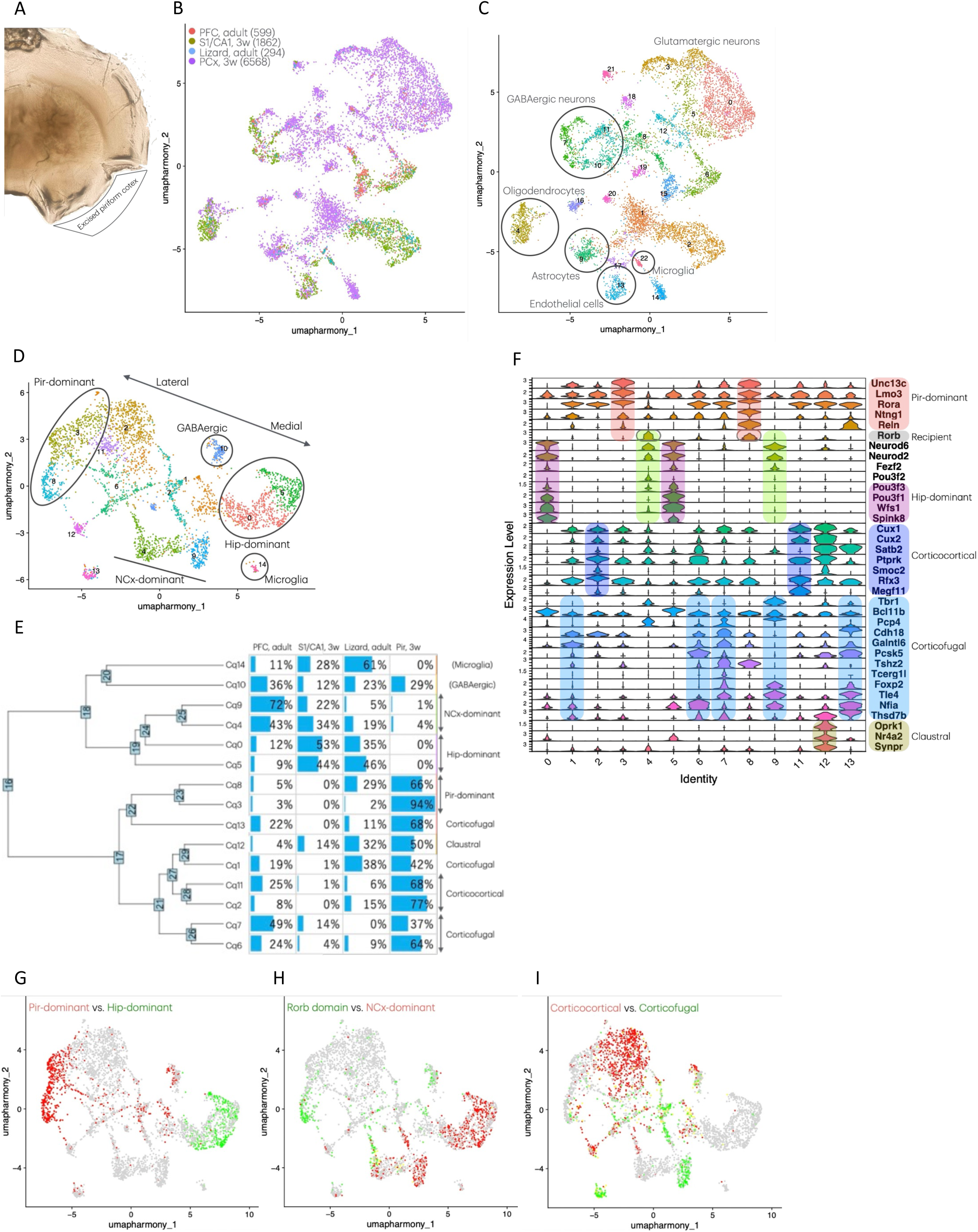
Inter-cortical and interspecies transcriptomic profiling of integrated datasets. (A) Representative excised tissue corresponding to the piriform cortex. (B) UMAP projection of harmony-integrated datasets colored by dataset origin. (C) UMAP projection colored by the Seurat clusters. The circles indicate clusters of GABAergic neurons, oligodendrocytes, astrocytes, endothelial cells, and microglia. (D) Clustering of glutamatergic neurons revealed pan-cortical neuronal populations and a mediolateral gradient distribution of area-specific neural clusters. (E) Left: Hierarchical clustering dendrogram of glutamatergic clusters. Right: distribution of cells derived from the adult PFC, 3-week S1/CA1, adult lizard cortex, and 3-week Pir, normalized for cell numbers across datasets. NCx-, Hip-, or Pir-dominant clusters were defined by the enrichment of cells from the PFC, S1/CA1, or Pir, respectively. Corticofugal, corticocortical, claustral clusters were annotated based on DEGs. (F) Violin plots showing the expression levels of DEGs and known molecular markers across Seurat clusters (GABAergic and microglial clusters excluded). Colored boxes indicate gene group characteristics of the Pir (red), Hip (purple), corticocortical neurons (indigo), corticofugal neurons (light blue), claustrum (brown), NCx (green), and sensory-recipient neurons (gray). (G–I) Distribution of cells with characteristic gene expression patterns in the UMAP space. (G) Pir-dominant (red) and Hip-dominant (green) expression signatures. (H) NCx-dominant expression (red) and Rorb domain expression (Rorb and related genes; green). (I) Corticocortical (red) and corticofugal (green) expression signatures.

Further differential gene expression analysis yielded 23 clusters (Supplementary Figure S2C,D). Ten Clusters were glial, including microglia (C4/7), oligodendrocytes (C9/22), astrocytes/pericytes (C10/11), oligodendrocyte progenitor cells (OPCs; C12/16), endothelial cells (C14), and vascular leptomeningeal cells (C19). Clusters C3/8/13/17 corresponded to GABAergic neurons, and the remaining nine corresponded to glutamatergic principal neurons. Although each Glu cluster exhibited distinct differentially expressed genes (DEGs)—including *Unc13c* for C2/6, *Ntng1* for C6, *Npsr1*, *Rspo2*, and *Satb2* for C5, *Tfap2d* for C15, *Grik3*, *Thsd7b*, *Cdh18*, and *Satb2* for C18, *Nr4a2*/*Nurr1*, *Oprk1*, *Ntng2*, and *Satb2* for C20, and *Pld5* and *Dach2* for C21—additional comparative analyses were required to better delineate the cellular architecture of the Pir (Supplementary Figure S2D).

### Interspecies data integration

To examine the tissue specificity of Pir neuronal architecture, we focused on neuronal lineage cells and incorporated three published RNA-seq datasets: two mouse NCx datasets, the adult prefrontal cortex (PFC) (Chen *et al*., 2022a), and the 3-week-old primary somatosensory cortex together with the hippocampal (Hip) CA1 region (S1/CA1) (Zeisel *et al*., 2015), and one dataset from the adult reptilian cortex (Tosches *et al*., 2018). These datasets enabled inter-cortical comparisons within mice as well as interspecies comparisons between mice and lizards. Together, we analyzed 9,323 neuronal lineage cells, defined by the expression of *Rbfox3/NeuN*, a pan-neuron marker, comprising 599 from adult PFC, 1,862 from 3-week-old S1/CA1, 294 from the reptilian cortex, and 6,568 from 3-week-old Pir. Harmony-based batch correction (harmony integration) successfully removed dataset-dependent clustering artifacts (Figure 1B). Subsequent dimensionality reduction revealed three GABA clusters and 16 Glu clusters (Figure 1C). A small fraction of *Rbfox3*^+^ cells fell into glial clusters, likely neuroblasts or immature neurons, and were removed from subsequent analyses. Glu and GABA populations were subsequently analyzed independently.

### Clustering of glutamatergic neurons

Using canonical markers (*Slc17a7*/*VgluT1*^+^ or *Slc17a6*/*VgluT2*^+^ for Glu and *Gad1*^+^ or *Gad2*^+^ for GABA), we identified 4,820 Glu and 1,194 GABA neurons. The extracted Glu population was subdivided into 15 glutamatergic clusters (Cq), with minor contamination by GABA neurons (Cq10) and Glia (Cq14) (Figure 1D; Supplementary Figure 3A). Cells from the four datasets were broadly distributed across the two-dimensional UMAP space, with each dataset displaying a distinct dispersion pattern that may reflect their developmental and evolutionary origins (Supplementary Figure 4A).

After normalizing the cell numbers for each dataset, we compared the cluster composition of the neurons (Figure 1E). Among the three mouse datasets, clusters Cq3/8 were predominantly composed of Pir-derived neurons (Pir-dominant), clusters Cq0/5 were enriched for S1/CA1-derived neurons (Hip-dominant), and clusters Cq4/9 were largely PFC-derived neurons (NCx-dominant). The identification of area-specific subtypes indicates the persistence of area-restricted neuronal programs (Tasic *et al*., 2018). Strikingly, Pir-dominant clusters were positioned in the upper-left region of the UMAP space, hippocampal clusters occupied the right region, and NCx-dominant clusters were interposed between them (Figure 1D). Their spatial arrangement in UMAP space mirrored the ontogenetic gradient along the mediolateral axis of the pallium: the PCx originated from the lateral pallium, the hippocampus from the medial pallium, and the NCx from the dorsal pallium, which lies between the two. Thus, transcriptomic diversity highlights the fundamental organizational principle of cortical development. Intermediate clusters (Cq1/2/6/7/11/12/13) were located in intermediate positions of the three Pir/NCx/Hip-dominant groups and likely represent pan-cortical glutamatergic lineages shared across regions (Figure 1D,E). The reptilian cortex-derived dataset contained a broad spectrum of cortical neurons, encompassing both pan-cortical and area-specific types, suggesting the evolutionary conservation of major glutamatergic neuron classes (Figure 1E).

To define the molecular identity of each cluster, we examined DEGs and known molecular markers (Figure 1F; Supplementary Figure 3A). Pir-dominant populations (Cq3/8) were defined by *Unc13c*, *Lmo3*, and *Rora*/*Nr1f1*, with *Ntng1*, *Reln*, *Rorb*/*Nr1f2*, and *Tshz2* being specifically enriched in Cq8 (Figure 1G; Supplementary Figure 5). Hip-dominant neurons (Cq0/5) were characterized by *Wfs1* and *Spink8* expression (Lein *et al*., 2007; Zeisel *et al*., 2015) and were further enriched in *Pou3f3/Brn1* and *Pou3f1/Oct6* (Figure 1G; Supplementary Figure 5). Marked expression of *Neurod6*/*Nex*/*Math2*, *Neurod2*, *Fezf2*, *Pou3f2/Brn2*, and *Tbr1* was observed in hippocampal and/or neocortical clusters, but was notably absent in Pir-dominant clusters (Figure 1H; Supplementary Figure 5). In the neocortex, *Rorb* and *Rora* mark spiny stellate neurons in layer IV that receive thalamic sensory inputs (Jabaudon *et al*., 2012; Vitalis *et al*., 2018). In particular, *Rorb*^+^ subtypes, including NCx-dominant Cq4 and PCx-dominant Cq8, may represent sensory-recipient neurons specialized for each sensory information processing (Zaremba *et al*., 2025). Consistently, the expression of putative *Rorb* target genes, *Thsd7a* and *Kcnk2*, was observed within both *Rorb* expression domains (Figure 1H; Supplementary Figure 5) (Clark *et al*., 2020; Zeppilli *et al*., 2025). Neocortical corticocortical projection neurons (Cq2/4/11) commonly expressed *Cux1*, *Cux2*, *Ptprk*, and *Rfx3*, with subsets also expressing *Satb2*, *Smoc2*, and *Megf11* (Figure 1I; Supplementary Figure 5). Neocortical corticofugal projection neurons (Cq1/6/7/9/13) showed a combination of high expression of *Tbr1*, *Bcl11b/Ctip2*, *Pcp4*, *Cdh18*, *Galntl6*, *Tshz2*, *Tcerg1l*, *Foxp2*, *Tle4*, *Nfia*, and *Thsd7b* (Figure 1I; Supplementary Figure 5). A distinct population (Cq12), marked by *Oprk1*, *Nr4a2/Nurr1*, and *Synpr*, corresponded to claustral neurons localized to the deepest region of the dorsally adjacent area to the piriform cortex and agranular insular cortex (AI) (Supplementary Figure 5) (Arimatsu *et al*., 2003; Zeisel *et al*., 2015; Puelles *et al*., 2016; Norimoto *et al*., 2020; Zeppilli *et al*., 2025). Collectively, we identified the specificity of Pir and the commonality of projection neurons across the cerebral cortex.

### In-depth characterization of paleocortical glutamatergic neurons

To further characterize the cellular composition of Pir glutamatergic neurons, we integrated our dataset with another published single-cell dataset derived from the adult Pir, subdivided into anterior and posterior halves (Zeppilli *et al*., 2025). A total of 6,103 glutamatergic neurons—including those from the adult anterior Pir (aaPir, 1,663 cells), adult posterior Pir (apPir, 2,275 cells), and 3-week-old Pir (3wPir, 2,165 cells)—were classified into 20 paleocortical clusters (Cps) characterized by distinct sets of DEGs. (Figure 2A; Supplementary Figure 3B,4B). Among these, DEG-based correspondences between Cq and Cp identified a variety of mutual markers, validating Pir-dominant neurons, corticocortical/upper layer (UL) neurons, corticofugal/deep layer (DL) neurons, and claustral neurons (Figure 2B; Supplementary Table 1). Pir-dominant clusters 1/4/5/12/17 were positioned next to the corticocortical UL clusters 0/2/3/11 and distant from the corticofugal DL clusters 8/10/15/16/19 in the UMAP space (Figure 2A). Next, we examined the RNA expression profiles of adult mice from the Allen Brain Atlas to provide the spatial expression patterns of the top 10 DEGs of each Cp (Lein *et al*., 2007) (Supplementary Figure 3B). Among Pir-specific DEGs, *Slc38a3*, *Ntng1*, *Rab3b*, *Tshz2*, *Igfbp5*, *Reln*, *Rorb*, *Whrn*, and *Pamr1* were selectively expressed in clusters Cp4/5/12/17 and neurons confined to the superficial zone of the piriform layer II (layer IIa), suggesting that Cp4/5/12/17 correspond to SL neurons and share sensory-recipient function with Cq8 spiny stellate neurons in the NCx (Figure 2C; Supplementary Figure 6A). DEGs of Cp1, such as *Unc13c*, *Rnf152*, *Plpp4*/*Ppapdc1a*, *Lmo3*, *Rora*, and *Foxo1*, and DEGs of Cp5/12, such as *Vav3*, *Nrsn2*, and *Tcerg1l*, were broadly expressed throughout layer II, although Tcerg1l expression was confined to the posterior Pir, encompassing both SL and Pyr neurons (Figure 2C; Supplementary Figure 6B) (Diodato *et al*., 2016; Zeppilli *et al*., 2025). Genes such as *Plk5*, *Penk*, *Stard8*, *Megf11*, *Ptprk*, *Tmem47*, *Lrmp*, *Tle1*, *Rfx3*, and *Rprm*, enriched in Cp0/2/3/7/18/19 and Cq2/11, were expressed in the upper layers (UL) in the NCx and restricted to the deeper zone of layer II (layer IIb) in the Pir, suggesting that NCx UL-corresponding neurons reside in Pir layer IIb (Figure 2C; Supplementary Figure 6C). *Pcsk5*, *Adcyap1*, *Cdh7*, and *Etv1*/*Er81* (the latter restricted to the posterior Pir), enriched in Cp10/15/19, were predominantly expressed in layer V in the NCx and limited to the superficial thin layer of Pir layer III (layer IIIa), indicating that Pir layer IIIa is equivalent to the NCx layer V (Figure 2C; Supplementary Figure 6D). Markers of the NCx layer VI, including *Tle4*, *Thsd7b*, and *Nfia*, enriched in Cp10/13, were located in the deeper layer III (layer IIIb) (Figure 2C; Supplementary Figure 6E). Other corticofugal neuron markers—*Pcp4*, *Galntl6*, *Pip5k1d*, *Cdh18*, and *Sema3e*—expressed in Cp8/10/13/15/16/19 were broadly distributed within the deep layers (DL) of the NCx and broad layer III of the Pir (Supplementary Figure 6F). Cp8/16 were characterized by *Slc17a6*/*VgluT2*, which was broadly expressed in layer III, along with the consistent expression of *Prkcq*, *Kctd8*, *Nxph1*, *Dock10*, *Cacna2d2*, *Meis1*, and *Foxp2* (Supplementary Figure 6G). The DEGs of Cp13—*Ccn2*/*Ctgf*, *Moxd1*, *Svil*, *Pcsk5*, *Tle4*, *Thsd7b*, *Cdh18*, *Nr4a2*/*Nurr1*, and *Cdh13*—formed a linear expression domain at the innermost portion of Pir layer III and NCx layer VIb/VII, demonstrating layer VIb/VII or subplate (SP)-equivalent neurons in the Pir (Figure 2C; Supplementary Figure 6D–F,H–J). DEGs of Cp14—*Oprk1*, *Cntnap3*, *Col11a1*, *Synpr*, and *Pld5*—were localized to the claustrum, and Cp6 DEGs—*Npsr1*, *Sema3e*, and *Spag16*—accumulated in the endopiriform nucleus, while *Nr4a2*/*Nurr1* and *Rspo2*, expressed in both Cp6 and Cp14, were found in these two nuclei as the claustral complex (Figure 6C; Supplementary Figure 6I). The DEGs of Cp11—*Smoc2*, *Cdh13*, *Trpc6*, *Stxbp6*, *Echdc2*, *Gpc3*, *Ano2*, *Nos1*, and *Cripsld1*—marked the dorsally adjacent agranular insular cortex (AI) (Supplementary Figure 6J). Cp9 neurons, mostly derived from apPir and not from 3wPir, expressed neural progenitor markers *Sox4* and *Sox11*, suggesting a progenitor-like state (Kaur *et al*., 2024; Zeppilli *et al*., 2025) (Figure 2B). Cp9 and other adult PCx-derived neurons (Cp7/17/18) exhibited high expression of mitochondrial genes, implying that they may originate from adult neural progenitors within the Pir, emerging during or after the gliogenic stage (Supplementary Figure 3B). Altogether, these analyses revealed a clear laminar organization in the piriform cortex (Figure 2D).

**Figure 2.**
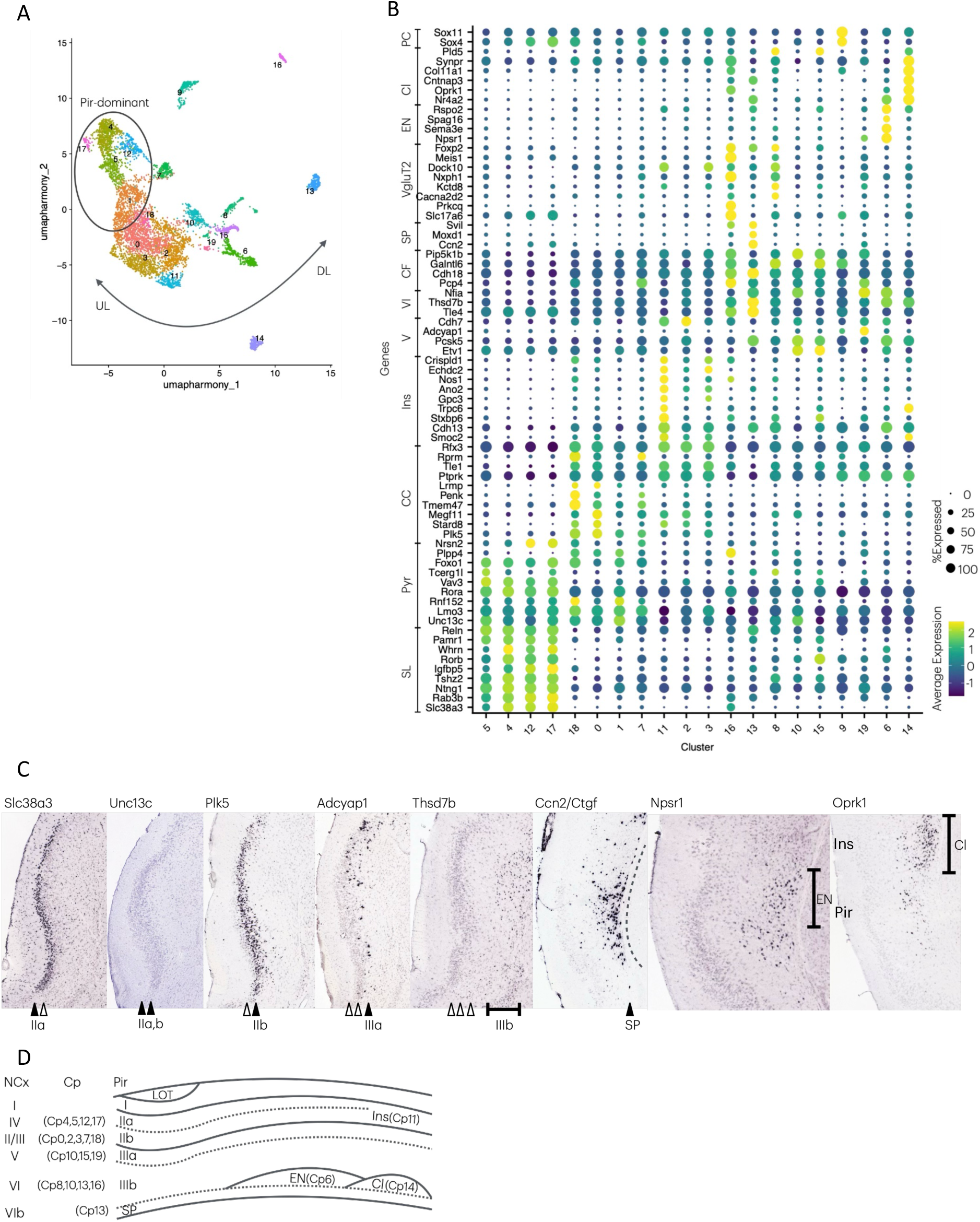
In-depth characterization of paleocortical glutamatergic neurons. (A) UMAP projection of 20 glutamatergic clusters derived from three Pir datasets (aaPir, apPir, and 3wPir) showing a gradient distribution of Pir-dominant neurons (circled), UL-type neurons, and DL-type neurons from left to right (two-headed curved arrow). (B) Dot plot showing the characteristic expression of DEGs. The dot size and color represent the proportion of cells expressing the gene and average expression level, respectively. (C) Spatial expression profiles of representative marker genes: Slc38a3 (layer IIa), Unc13c (layers IIa and IIb), Plk5 (layer IIb), Adcyap1 (layer IIIa), Thsd7b (layer IIIb), Ccn2/Ctgf (SP), Npsr1 (EN), and Oprk1 (Cl). Filled and open arrowheads indicate layers with and without detectable expression, respectively. The dotted line indicates the boundary beneath the innermost portion of layer III. (D) Schematic model of neocortex-like laminar organization in the piriform cortex. The Pir layers and sublayers are indicated by solid and dotted lines, respectively. The corresponding NCx layers are shown on the left. Clusters (Cp) with characteristic spatial expressions were mapped to putative laminar locations. Abbreviations: CC, corticocortical; CF, corticofugal; Cl, claustrum; EN, endopiriform nucleus; Ins, insular; LOT, lateral olfactory tract; PC, progenitor cells; Pyr, superficial pyramidal; SL, semilunar; SP, subplate.

According to previous birthdate analyses, Pir neurons are generated in the temporal order of layer III, layer IIa, and layer IIb, reported as an inter-layer “inside-out” and an intra-layer “outside-in” mechanism (Martin-Lopez *et al*., 2019; Baumann *et al*., 2025), which is partially different from the NCx, where later-born neurons migrate past earlier-born populations to form an inside-out structure (Frantz and McConnell, 1996; Hevner *et al*., 2003; Hevner *et al*., 2006; Shen *et al*., 2006; Molyneaux *et al*., 2007). We found that Pir layers III, IIa, and IIb were composed of corticofugal projection neurons, sensory-recipient neurons, and corticocortical neurons, respectively, and corresponded to NCx layers V/VI, IV, and II/III. Although late-born Pir corticocortical neurons remain beneath sensory-recipient neurons, the Pir adopts a temporal pattern analogous to NCx. Another difference between the NCx and Pir is the cellular composition of corticocortical (UL) and corticofugal (DL) neurons. The UL/DL ratio of the adult PFC was 3:5, whereas that of the 3wPir was 3:2, indicating that UL neurons in the Pir shared a large fraction (Figure 2A; Supplementary Figure 3B). Consistently, NCx-dominant Cq9 was observed in DL neurons, suggesting a functional constraint on the tissue architecture in the cerebral cortex (Figure 1F). NCx layer markers have contributed to identifying the cellular composition of the entire cortex outside the NCx and reconstructing pallial evolution in vertebrates (Molyneaux *et al*., 2007; Suzuki and Hirata, 2013; Luzzati, 2015; Klingler, 2017). Unfortunately, because the regulatory logic for determining cell identity is evolutionarily plastic and developmentally dynamic, the whole picture remains ambiguous (Nomura *et al*., 2018; Nasu *et al*., 2020; Tosches, 2021). In this study, we identified universal molecular markers enabling comparative analyses across the NCx and Pir regions. Taken together, the glutamatergic neuron landscape reflects both conserved developmental trajectories shared across cortical regions and specialized programs that define area-specific identity. Transcriptomic analysis with spatial information revealed NCx-like layer formation in the Pir, with an inverted positioning of sensory-recipient and corticocortical neuron compartments relative to the NCx.

### Clustering of GABAergic neurons

Across Pir, PFC, S1/CA1, and reptilian cortex, the GABAergic population segregated into 12 clusters of GABAergic neurons (Cg), seven of which represented bona fide interneuron populations (Figure 3A; Supplementary Figure 3C). GABAergic neurons from all four datasets were primarily distributed in Cg1/2/3/4/5/11 (Supplementary Figure 4C). After a reliable estimation to exclude contamination in the clustering, we found 2,332 glutamatergic and 447 GABAergic neurons in the Pir and 401 glutamatergic and 79 GABAergic neurons in the PFC. The excitatory/inhibitory (E/I) ratio in the mouse Pir was 5.2:1, which was comparable to that in the PFC and just higher than the widely accepted ratio of 4:1 in the mouse NCx.

**Figure 3.**
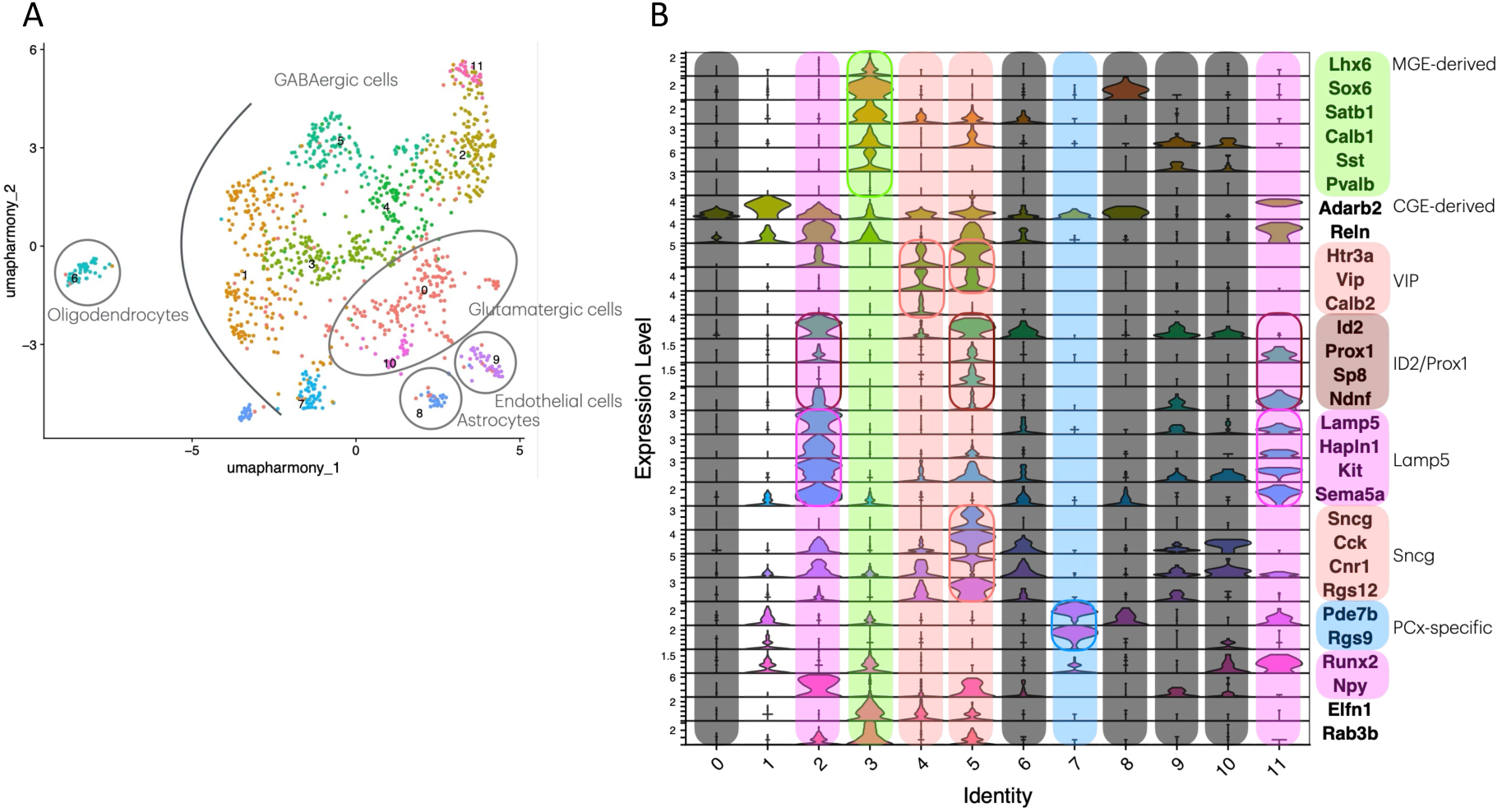
Clustering of GABAergic neurons. (A) UMAP projection of GABAergic neurons from the harmony-integrated datasets. Clusters enclosed in circles represent contaminated non-GABAergic populations, including glutamatergic neurons, astrocytes, oligodendrocytes, and endothelial cells. (B) Violin plots showing the expression levels of the DEGs and established molecular markers. Colored boxes indicate the genes characteristic of specific GABAergic subtypes.

Two major developmental lineages have been identified: MGE-derived and CGE-derived neurons (Tasic *et al*., 2018; Wang *et al*., 2022; Machold and Rudy, 2024). MGE-derived neurons were grouped predominantly within cluster Cg3, which expressed the canonical markers *Lhx6*, *Sox6*, *Satb1*, and *Calb1*/*Calbindin*/*CB* (Figure 3B; Supplementary Figure 4C). This cluster included both *somatostatin* (*SST*)-positive and *parvalbumin* (*PV*)-positive interneurons, which represent the major MGE-derived subtypes. In contrast, clusters Cg1/2/4/5/11 were characteristic of CGE-derived neurons, as defined by the expression of *Adarb2*. Among them, Cg2/4/5/11 were further marked by CGE-enriched genes such as *Id2*, *Prox1*, and/or *Sp8*, while Cg1 lacked distinct DEGs, suggesting that Cg1 may represent immature or transitional GABAergic neurons. Within the CGE lineage, clusters Cg4/5 were classified as *Htr3a*^+^*VIP*^+^ neurons, although not all cells shared the expression of *Htr3a* and/or *Vip*. Cluster Cg4 contained *Calb2*/*Calretinin*/*CR*^+^ neurons, while Cg5 corresponded to *Sncg*^+^ neurons characterized by high expression of *Sncg*, *Ccr*, *Cnr1*, and *Rgs12*. Clusters Cg2/11 corresponded to *Lamp5*^+^ neurons that expressed high levels of *Lamp5*, *Hapln1*, *Kit*, and *Sema5a*. Cluster Cg11 was enriched in the Pir-derived dataset and composed of *Lamp5*^+^*Ndnf*^+^ neurons, showing high expression of *Ndnf* and *Runx2*. In contrast, Cg2 neurons were *Lamp5*^+^*Npy*^+^ neurons that expressed *Npy* and exhibited stronger *Lamp5* expression than Cg11 neurons. Cluster Cg7 consists exclusively of Pir-derived neurons characterized by the expression of *Rgs9* and *Pde7b* (Thomas *et al*., 1998; Reyes-Irisarri *et al*., 2005). According to the Allen Brain Atlas, both genes show low expression levels in the Pir but are highly expressed in the olfactory tubercle (OT) and striatum (Str)—regions that are ventrally adjacent to the Pir— and are virtually absent in the NCx (Supplementary Figure 6K) (Lein *et al*., 2007). This expression pattern suggests that the *Rgs9*^+^*Pde7b*^+^ GABAergic subtype identified in the Pir may not share a common origin with neocortical GABAergic neurons. Instead, these cells may represent a lineage more closely related to the neurons of the OT and Str, which are derived from the LGE (De Carlos *et al*., 1996; Wichterle *et al*., 2001; Legaz *et al*., 2005; García-Moreno *et al*., 2008; Klingler, 2017). Together, these transcriptional profiles suggest that the GABAergic neurons in the mouse paleocortex retain the classical dual lineage organization (MGE vs. CGE) observed in the neocortex but also exhibit a minor GABAergic population potentially originating from the LGE, pointing to a region-specific diversification of inhibitory circuit composition in the paleocortex.

### Both Glutamatergic and GABAergic neurons contribute to the area-selective vulnerability

To explore the functional implications of the identified Glu and GABA clusters, we performed disease ontology enrichment analysis. DEGs identified in the Hip-dominant glutamatergic clusters Cq0/5 were enriched for genes implicated in Alzheimer’s disease, whereas those in the NCx-dominant clusters Cq4/9 showed significant enrichment for genes associated with toxic encephalopathy and Pick’s disease (Figure 4A). Cq0/5 and Cq4/9 also shared a subset of DEGs related to synucleinopathies, including Parkinson’s disease (PD). In contrast, Pir-dominant clusters Cq3/8 were enriched for DEGs associated with autism spectrum disorder (ASD) and intellectual disability. These DEGs were also observed in the pan-cortical clusters Cq1/2/6/7/11/13 but not in Cq4/9 or Cq0/5, indicating a distinct transcriptional signature of the area-specific lineage as well as a common functional circuit across areas. Some abnormalities in functional circuits in the cerebral cortex are causally linked to clinical symptoms and range from the neocortical to the paleocortical areas.

**Figure 4.**
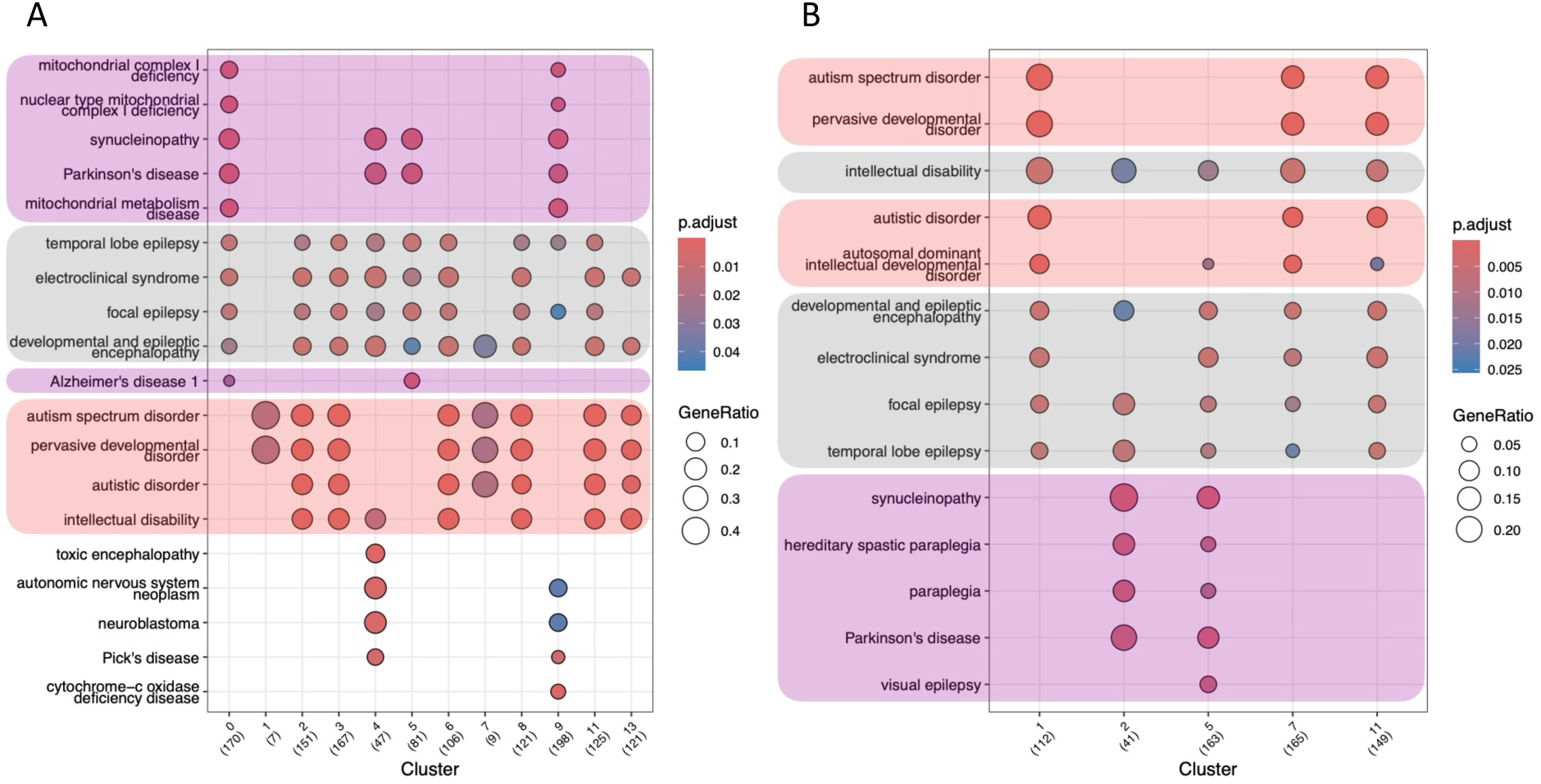
Gene enrichment analysis. (A) Dot plot showing disease-associated gene enrichment in glutamatergic clusters (Cq). (B) Dot plot showing disease-associated gene enrichment for GABAergic clusters (Cg). The dot size and color represent the proportion of cells expressing the gene set and enrichment score, respectively. Colored boxes indicate cortical area specificity: pan-cortical (gray), Hip-dominant (purple), Pir-dominant, and NCx-dominant (red). The number of associated genes in each cluster is indicated in parentheses.

A comparable pattern of area-specific disease association was observed among GABAergic clusters (Figure 4B). Approximately 77.1% (209/271) of cells belonging to clusters Cg2/5 were derived from S1/CA1 or PFC samples, and their DEGs were predominantly linked to synucleinopathy and Parkinson’s disease. Conversely, 94.9% (261/275) of the cells in clusters Cg1/7/11 originated from Pir samples and exhibited enrichment for DEGs associated with ASD and intellectual developmental disorders. Both glutamatergic and GABAergic neurons retained independent risk factors for epilepsy across cortical areas. These results suggest that transcriptional specialization among cortical regions is mirrored by their distinct susceptibilities to neurological diseases. Notably, both glutamatergic and GABAergic neurons contribute to these region-specific disease associations in a similar pattern, implying that not only the developmental patterning of cortical subtypes but also region-dependent circuit organization, particularly the local integration between excitatory principal neurons and inhibitory interneurons, may underlie the selective vulnerability characteristic of diverse neurological and neurodevelopmental disorders.

## Discussion

### Conserved developmental programs across mammalian cortices

In this study, we performed single-cell transcriptomic profiling of Pir, combined with inter-cortical and interspecies comparative analyses, to delineate the transcriptomic landscape of cortical neurons. We identified neuronal subtypes shared among the Pir, NCx, and hippocampus, suggesting that subsets of glutamatergic neurons exhibit conserved transcriptomic features across cortical regions, indicating a pan-cortical lineage underlying the fundamental excitatory circuitry. Meanwhile, their gradient distribution strongly supports the notion that the transcriptional heterogeneity of cortical neurons retains a molecular imprint of their embryonic origin. From a developmental perspective, this organization suggests that the molecular programs specifying neuronal identity are not completely homogenized during postnatal maturation but instead preserve positional information established during pallial patterning. This principle resonates with classical neurodevelopmental models proposing that the pallium generates diverse cortical areas through graded signaling along its mediolateral axis (Marín and Rubenstein, 2001). Our transcriptomic data provide molecular evidence supporting this model at the single-cell resolution.

### A six-layered organizational framework in the paleocortex

Classically, the PCx, including the Pir, has been characterized by a three-layered architecture with sublayers and has therefore been regarded as evolutionarily distinct from other cortices, representing a cortex closer to an ancestral form shared with the reptilian cortex, which also exhibits a three-layered organization comprising DL-type and UL-type neurons. To clarify the evolutionary position of the PCx, we compared the cytoarchitecture of the mammalian NCx, PCx, Hippocampus, and the reptilian cortex. In this study, we identified a deep-layer organization in the Pir comparable to NCx DL-type corticofugal neurons. In the superficial Pir layers, we detected both an NCx UL-type corticocortical neuronal layer and a sensory recipient neuronal layer. Each neuronal population is also found in the whole pallium of reptiles, suggesting the existence of evolutionarily conserved cell types that predate the diversification of mammalian cortex.

Spatial transcriptomic analyses further revealed laminar segregation in the Pir corresponding to NCx layers II/III, IV, V, VI, and VIb, although the layer II/III-like compartment lacked distinct histological separation. The distribution of Pir UL-type and DL-type corticofugal neurons followed a pattern reminiscent of the NCx, residing in the superficial and deeper layers, respectively. Intriguingly, the relative positioning of DL-type and UL-type (or sensory recipient) neurons is not conserved even among reptiles; for instance, they reside in layers IIa and IIb in turtles, but are reversed in alligators (Dugas-Ford *et al*., 2012; Tosches *et al*., 2018; Briscoe and Ragsdale, 2018). Furthermore, while Pir layer III is abundant in DL-type neurons, it is sparse in reptilian layer III (Nomura *et al*., 2013, J. Exp. Zoo.). These observations suggest that the specific laminar organization of UL-type- and DL-type neurons is a characteristic feature of the mammalian cortex. In contrast, its prevalence among amniotes remains unclear.

Based on previous birthdate analyses, we propose that the Pir follows an NCx-like sequential neurogenesis, even though its corticocortical and sensory-recipient neurons occupy inverted positions relative to the NCx (Martin-Lopez *et al*., 2019; Baumann *et al*., 2025). A key distinction exists in their afferent connectivity: NCx layer IV neurons receive thalamocortical inputs ascending from the apical side of the cortical plate, whereas PCx neurons receive olfactory inputs via the lateral olfactory tract (LOT) along the cortical surface. This difference may influence the migration behavior or maturation sites of sensory-recipient neurons. The precise mechanisms underlying the inverted arrangement of the PCx remain subjects for future investigations. Together, our findings support a revised model in which the paleocortex represents not an immature three-layered allocortex but a divergent specialization of a six-layered isocortex.

### Area-restricted developmental programs

In addition to conserved features, we identified area-restricted developmental programs that shape Pir-and Hip-specific neuronal subtypes. Our analyses revealed two Pir-dominant glutamatergic subtypes and one Pir-specific GABAergic subtype, collectively referred to as paleocortical neurons (PCx neurons), suggesting that they may represent the evolutionary innovations underlying the paleocortex-specific circuitry. Nevertheless, low-abundance homologous populations of reptiles cannot be excluded.

One subtype of PCx glutamatergic neurons, corresponding to *Rorb*^+^ SL cells, functions as an olfactory sensory recipient. In the NCx, *Rorb*^+^ spiny stellate neurons in layer IV are well-established thalamocortical sensory recipient neurons. In the Pir, the expression of the paralogous gene *Rora* was observed in recipient neurons, which is shared with sensory recipient neurons in the NCx, birds, and reptiles (Briscoe and Ragsdale, 2018). The expression of *Rorb* in the primary somatosensory cortex is critical for fate specification and the formation of thalamocortical sensory inputs (Jabaudon *et al*., 2012; Clark *et al*., 2020; Katayama *et al*., 2024). Moreover, a distinct molecular identity of sensory recipient neurons has recently been reported in the human primary visual cortex (Qian *et al*., 2025). Collectively, although sensory recipient neuron populations segregate into distinct transcriptomic clusters across cortical regions, it remains possible that these populations share a common developmental program that generates a conserved sensory-recipient archetype, with subsequent regional diversification producing transcriptomic and functional differences. Further developmental and phylogenetic analyses are required to test this hypothesis.

The Pir is a distinctive cortical region containing LGE-derived GABAergic populations (De Carlos *et al*., 1996; Wichterle *et al*., 2001; Legaz *et al*., 2005; García-Moreno *et al*., 2008; Klingler, 2017). A recent study reported the presence of LGE-derived GABAergic neurons in the avian pallium (Zeremba *et al*., 2025). Consistent with these observations, our analyses revealed a minor GABAergic population in the PCx that potentially originates from the LGE, in addition to the canonical MGE- and CGE-derived interneurons. This PCx-specific GABAergic subtype may contribute to PCx-specific circuit function. An important question for future studies is whether the incorporation of LGE-derived GABAergic neurons into the PCx reflects an ancestral pattern of inhibitory neuron distribution that predates the strict MGE/CGE segregation characteristic of the neocortex or represents a mammalian PCx-specific specialization. Given the limited sampling of reptilian cortical datasets, additional comparative and developmental analyses are required to distinguish between these evolutionary scenarios. By delineating paleocortical neuronal populations at the molecular level, this study provides new insights into how embryonic pallial patterning gives rise to the evolutionary and functional diversity of conserved, multilayered cortical architectures.

### Disease vulnerability

The Pir has historically been overlooked as a primary site of neurological dysfunction, largely because of the lack of universal markers for comparative analysis across the NCx and PCx. The identification of these markers enabled gene enrichment analyses, which revealed disease associations specific to the cortical areas. Pir-dominant and pan-cortical clusters were enriched for DEGs associated with Autism Spectrum Disorder (ASD) and intellectual disability, suggesting the existence of shared functional circuits spanning the neocortical and paleocortical regions. Recent clinical and experimental evidence has increasingly linked altered olfaction to ASD patients and mouse models (Boudjarane *et al*., 2017; Sweigert *et al*., 2020; Geramita *et al*., 2020). Histological abnormalities in the Pir may represent autism-related phenotypes (Mihalj *et al*., 2024). Notably, both glutamatergic and GABAergic neurons exhibited similar patterns of region-specific disease associations, indicating that coordinated alterations in excitatory and inhibitory (E/I) circuits may underlie the pathogenesis of neurodevelopmental disorders. Together, the transcriptomic landscape of cortical neurons defined here provides a molecular framework for understanding region-specific disease susceptibility and offers insights into the evolutionary diversification of cortical circuits in mammals.

## Materials and Methods

### Tissue Preparation, Nuclei isolation, and 10X single nuclei RNA-seq

A pair of male C57BL/6N mice (postnatal day 23–25; CLEA Japan) were anesthetized with propofol (2,6-diisopropylphenol; 10 mg/kg) via retro-orbital injection and transcardially perfused through the left ventricle with Hank’s Balanced Salt Solution (HBSS). The piriform cortex from both hemispheres (Bregma +1.0 mm to −1.0 mm) was microdissected, including 1 mm of both anterior and posterior subdivisions (Figure 1A; Supplementary Figure 1A). Tissues from two animals were minced and pooled prior to further processing. Single cells and nuclei were isolated using the Papain Dissociation System (Worthington Biochemical Corp.) and Single Nucleus Isolation Kit for Neuronal Tissues/Cells (Invent Biotechnologies Inc.), respectively, according to the manufacturers’ protocols. Nuclear integrity and yield were assessed by Trypan Blue staining and quantified using a hemocytometer (Supplementary Figure 1B). Nuclei were adjusted to a concentration of 900 nuclei/µL and processed for library preparation for next-generation sequencing (NGS) using the Chromium Next GEM Single Cell 3’ Kit v3.1 on a Chromium Controller (10x Genomics), following the manufacturer’s instructions. Libraries incorporating unique molecular identifiers (UMIs) were sequenced using an Illumina NovaSeq platform. All animal experiments were conducted in accordance with the institutional (Kumamoto University) guidelines and were approved by the Institutional Animal Care and Use Committee of Kumamoto University.

### Read processing and quality control

Raw sequencing data were processed using Cell Ranger software (v7; 10x Genomics) to align the reference mouse genome (mm10/GRCm38), count unique molecular identifiers (UMIss), and perform initial quality control. To minimize technical noise and unwanted biological variations, we performed sctransform-based normalization with Seurat (v4.3; Satija *et al*., 2015, Hao *et al*., 2021) and removed variables related to the sequencing depth per cell (nCount_RNA) and mitochondrial gene fraction (percent.mt). Normalized data were processed using Principal Component Analysis (PCA) for dimensionality reduction and the Uniform Manifold Approximation and Projection (UMAP) algorithm (McInnes *et al*., 2018) with 30 nearest neighbors to visualize cells in a two-dimensional space. Cell clusters were identified based on a shared nearest-neighbor graph (resolution = 0.8).

To find differentially expressed genes (DEGs) among clusters, cut-off values were set as follows: the expression frequency within the cluster was 25% or more (min.pct = 0.25), and the fold change of expression across clusters was 1.19-fold or more (logfc.threshold = 0.25). A phylogenetic tree was constructed based on a distance matrix of the average gene expression in each cluster. Relative gene expression across clusters or batches was visualized in a heatmap constructed using highly changed DEGs. Particular cell types were filtered by specific gene expression as follows: neuronal lineages (*Rbfox3* > 0.1, *Aqp4* < 1, *Sox10* <1, and *Cx3cr* < 1), glutamatergic neurons (*Slc17a7* > 0.1 or *Slc17a6* > 1), and GABAergic neurons (*Gad1* > 1 or *Gad2* > 1).

### Data integration

The following public data were used in this study: adult prefrontal cortex (Chen *et al*., 2022a), 3-week-old primary somatosensory cortex and hippocampal CA1 region (Zeisel *et al*., 2015), adult lizard (Pogona vitticeps) cortex (Tosches *et al*., 2018), and adult anterior and posterior piriform cortices (Zeppilli *et al*., 2025). The lizard dataset was mapped to the reference genome (genome assembly pvi1.1) using STARsolo (Kaminow *et al*., 2021). Data integration and batch correction were achieved using harmony-based batch correction (harmony integration) in a Seurat pipeline (Korsunsky *et al*., 2019).

### Gene Enrichment Analysis

Gene Enrichment Analysis was performed using clusterProfiler (Yu *et al*., 2012). Common DEGs shared by 90% or more cells in each cluster were compared with disease-associated gene lists from the Disease Ontology (Baron *et al*., 2026).

### Database-based spatial expression profiling

Spatial gene expression patterns were examined using in situ hybridization (ISH) data from adult mice from the Allen Mouse Brain Atlas (Lein *et al*., 2007). Layer- and region-specific expression was manually curated based on coronal section images and annotated anatomical boundaries, including cortical layers IIa, IIb, IIIa, IIIb, subplate, claustral complex, and insular cortex.

### Statistical analysis

All statistical analyses were conducted in R. Unless otherwise specified, statistical significance was defined as an adjusted p-value of < 0.05. Data visualization was performed using the Seurat and ggplot2 software.

## Data availability

The original RNA-seq data presented in this study are publicly available from the Gene Expression Omnibus (GEO accession number: GSE333337).

## Acknowledgement

We thank Keiichiro Yasunaga for technical support and Novogene Co., Ltd. for assistance with the next-generation sequencing. This work was supported by JSPS KAKENHI Grant Number JP15K18957 (MN) and the MEXT-Supported Program for the Inter-University Research Network for High Depth Omics, IMEG, Kumamoto University. This manuscript was drafted using ChatGPT (GPT-4), Gemini, and Paperpal and refined by the authors. The authors declare no conflict of interest.

## Author contributions

M.N. conceived and designed the research, performed all experiments and analyses, and wrote the manuscript. E.S., T.W., and K.S. helped performing the experiments. All authors edited and approved the final version of the manuscript.

