## Supplementary information for "A conserved lamination pattern in the paleocortex and the neocortex revealed by single-cell RNA analyses"

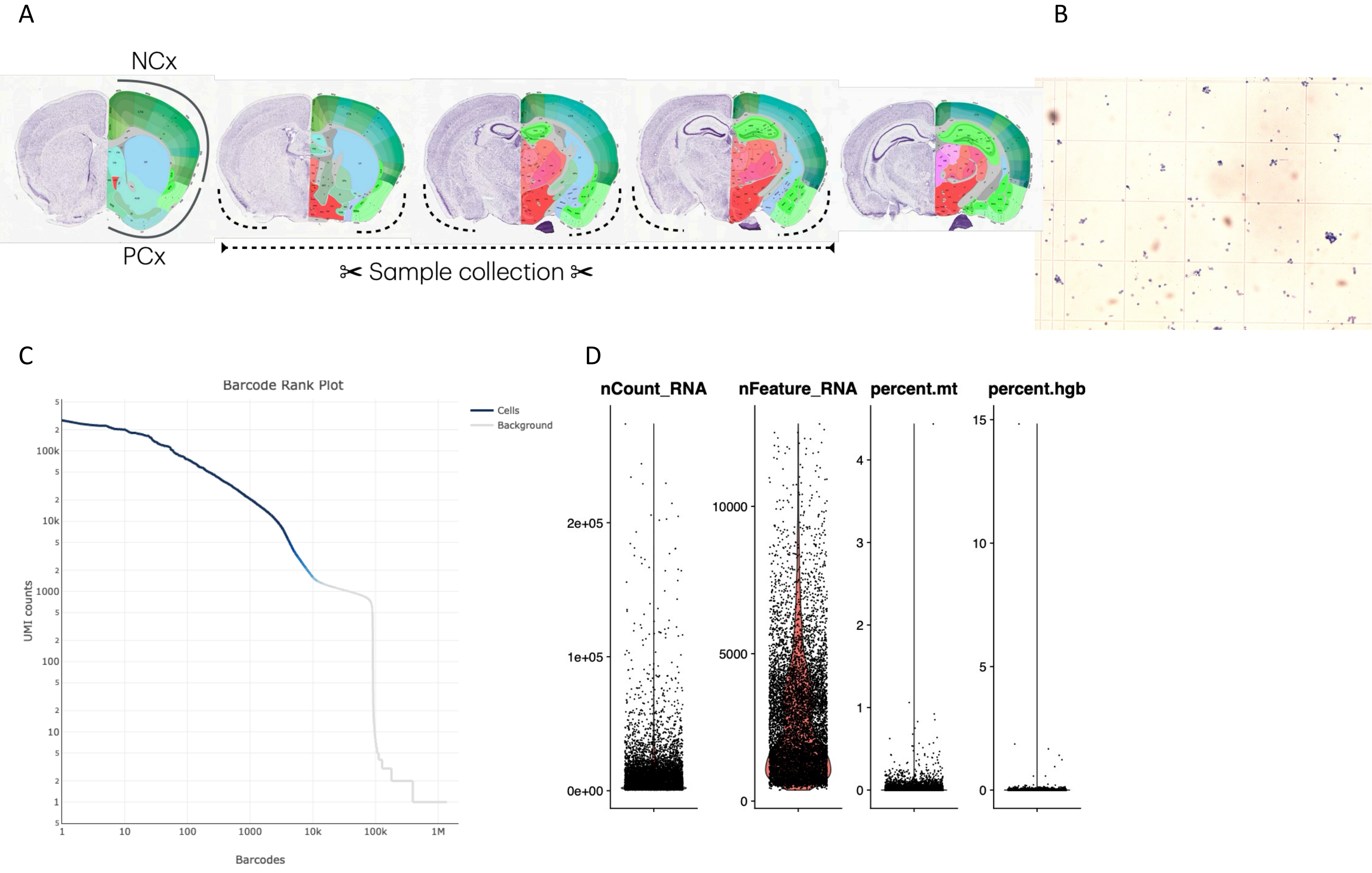


Supplementary Figure 1. Sample collection and quality control.
(A) Schematic illustration of the sample-collection strategy. Dotted lines indicate the piriform cortex targeted for transcriptomic profiling in this study. (B) Quantification of nuclei stained with Trypan Blue following single-nucleus dissociation. (C) Barcode rank plot showing the distribution of UMI counts per cell barcode, indicating high-quality library preparation. (D) Violin plots showing the sequencing depth per cell (nCount_RNA), number of detected genes (nFeature_RNA), mitochondrial gene fraction (percent.mt), and red blood cell contamination (percent.hgb).

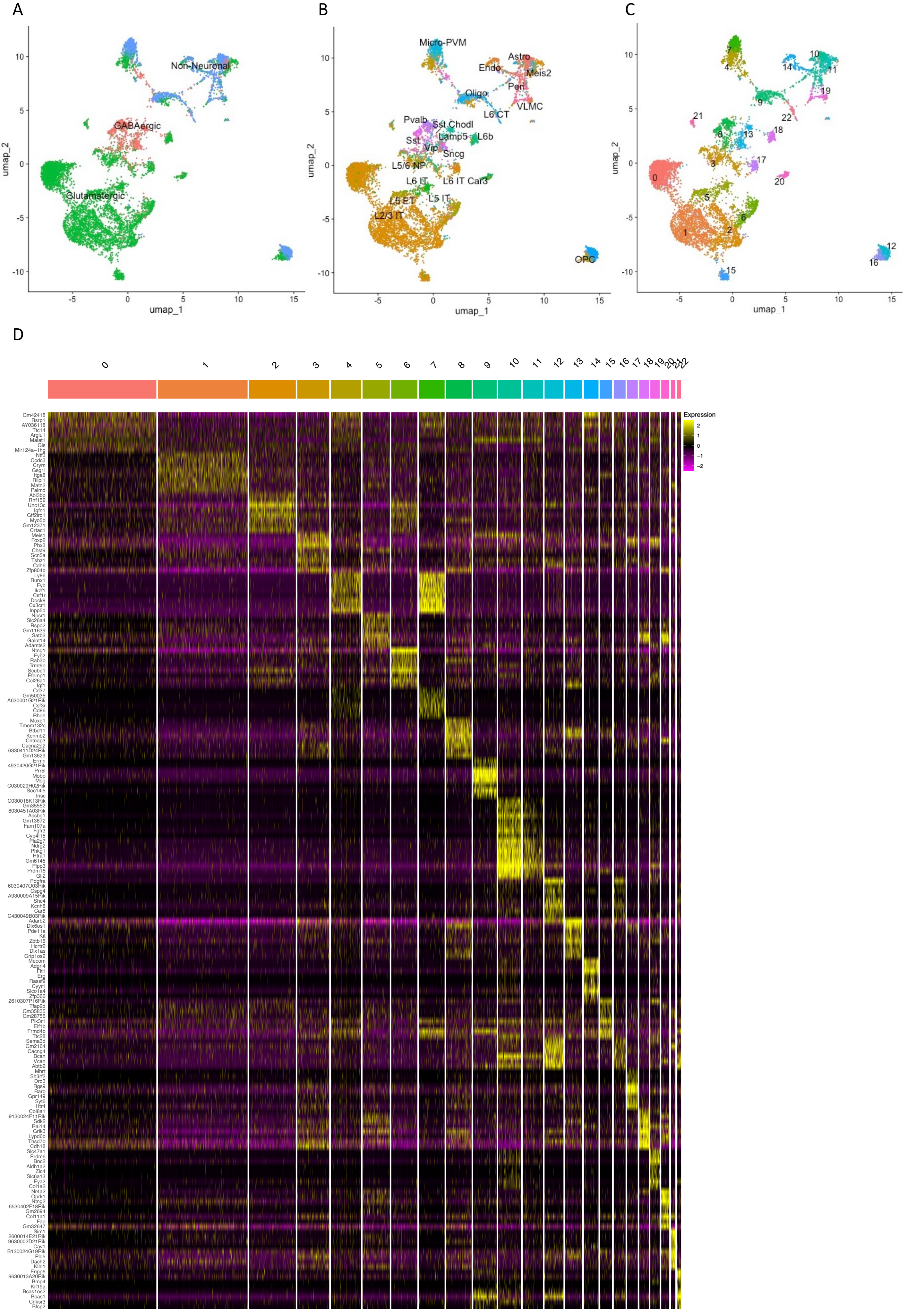
A
B
C
D
umap_2
umap_1
Non-Neuronal
GABAergic
Glutamatergic
Micro-PVM
Astro
Endo
Meis2
Peri
Oligo
VLMC
L6 CT
Pvalb
Sst Chodl
Lamp5
L6b
Sst
Vip
Sncg
L5/6 NP
L6 IT
L6 IT Car3
L5 ET
L5 IT
L2/3 IT
OPC
Expression
Gm42418
Rsrp1
AY036118
Ttc14
Arglu1
Malat1
Gls
Mir124a−1hg
Ntf3
Ccdc3
Crym
Gsg1l
Itga8
Rilpl1
Matn2
Palmd
Abi3bp
Rnf152
Unc13c
Igfn1
Gtf2ird1
Myo5b
Gm12371
Crtac1
Meis1
Foxp2
Pbx3
Chst9
Scn5a
Tshz1
Cdh6
Zfp804b
Ly86
Runx1
Fyb
Ikzf1
Csf1r
Dock8
Cx3cr1
Inpp5d
Npsr1
Slc26a4
Rspo2
Gm11639
Satb2
Galnt14
Adamts2
Ntng1
Fyb2
Rab3b
Trmt9b
Scube1
Efemp1
Col26a1
Igf1
Cd37
Gm50035
A630001G21Rik
Csf3r
Cd86
Rhoh
Moxd1
Tmem132c
Btbd11
Kcnmb2
Cntnap3
Cacna2d2
6330411D24Rik
Gm13629
Ermn
4930420G21Rik
Prr5l
Mobp
Mog
C030029H02Rik
Sec14l5
Insc
C030018K13Rik
Gm35552
8030451A03Rik
Acsbg1
Gm13872
Fam107a
Fgfr3
Cyp4f15
Pla2g7
Ndrg2
Phkg1
Htra1
Gm6145
Plpp3
Prdm16
Gli2
Pdgfra
6030407O03Rik
Cspg4
A930009A15Rik
Shc4
Kcnh8
Car8
C430049B03Rik
Adarb2
Dlx6os1
Pde11a
Kit
Zbtb16
Hcrtr2
Dlx1as
Grip1os2
Mecom
Adgrl4
Flt1
Erg
Rassf9
Cyyr1
Slco1a4
Zfp366
2610307P16Rik
Tfap2d
Gm35835
Gm28756
Pik3r1
Eif1b
Frmd4b
Ttc28
Sema3d
Gm2164
Cacng4
Bcan
Vcan
Abtb2
Mhrt
Sh3rf2
Drd3
Rgs9
Rarb
Gpr149
Syt6
Htr4
Col8a1
9130024F11Rik
Sdk2
Rai14
Grik3
Lypd6b
Thsd7b
Cdh18
Slc47a1
Prdm6
Bnc2
Aldh1a2
Zic4
Slc6a13
Eya2
Col1a2
Nr4a2
Oprk1
Ntng2
6530402F18Rik
Gm2694
Col11a1
Fap
Gm32647
Sim1
2600014E21Rik
9630002D21Rik
Cav1
B130024G19Rik
Pld5
Dach2
Klhl1
Enpp6
9630013A20Rik
Bmp4
Kif19a
Bcas1os2
Bcas1
Cnksr3
Bfsp2

Supplementary Figure 2. Transcriptomic profiling of a single dataset generated in this study.

(A) UMAP projection classifying cells into glutamatergic (green), GABAergic (red), and non-neuronal populations (blue). (B) UMAP projection illustrating the subclasses based on the cortical lineage identity. (C) UMAP projection colored by Seurat clusters (0-22). (D) Heatmap showing the top eight DEGs for each Seurat cluster. The rows and columns represent genes and individual cells, respectively. Expression levels are indicated by yellow for high and magenta for low expression.

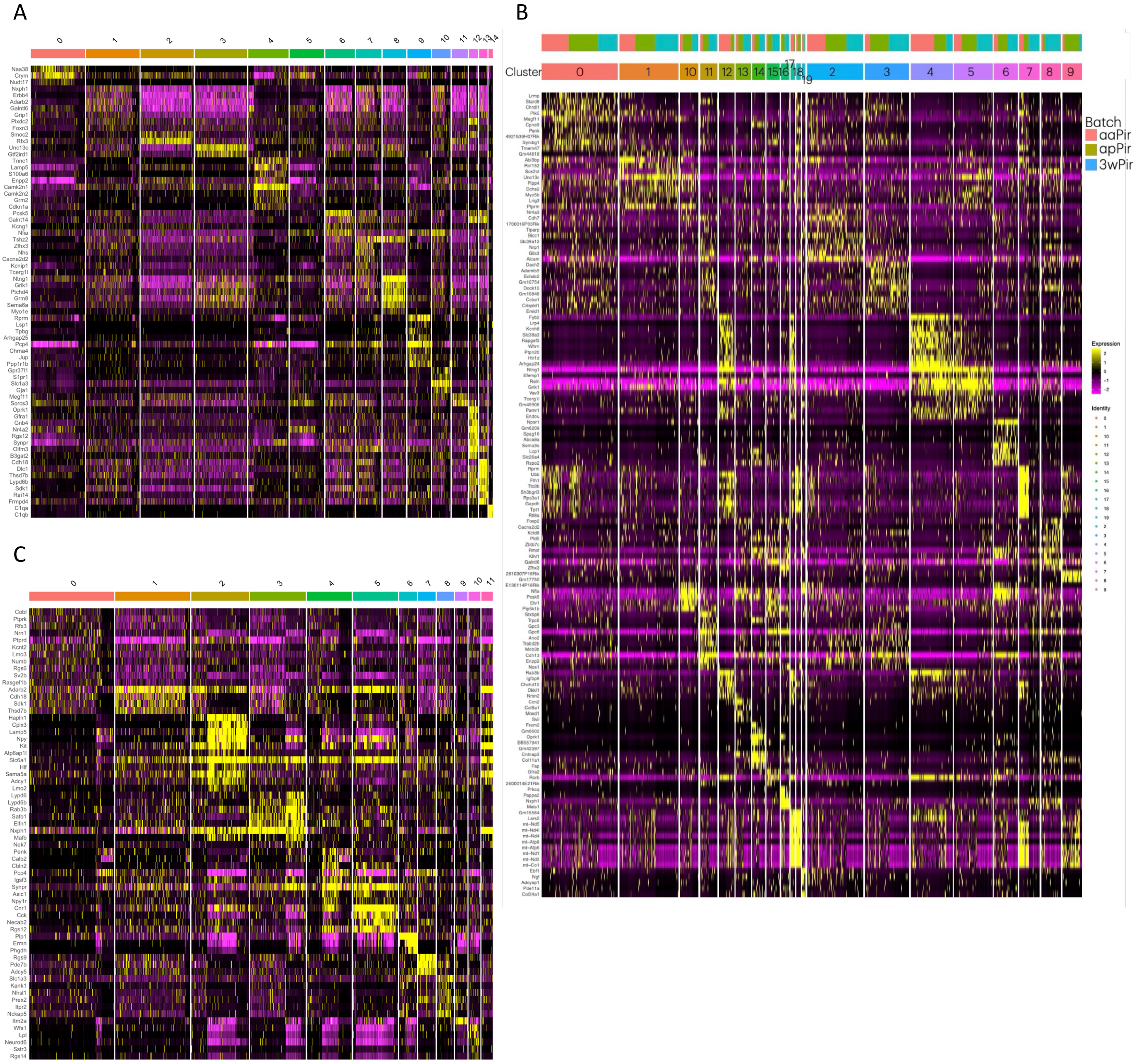


Supplementary Figure 3. DEG-based clustering of the integrated datasets.
(A) Heatmap of glutamatergic clusters (Cq0-14) based on the top 20 DEGs for each Seurat cluster across four inter-cortical and interspecies datasets. (B) Heatmap of paleocortical glutamatergic clusters (Cp0-19) based on the top seven DEGs for each Seurat cluster across three piriform cortex datasets (aaPir, apPir, and 3wPir). (C) Heatmap of GABAergic clusters (Cg0-11) based on the top 20 DEGs for each Seurat cluster across four inter-cortical and interspecies datasets. Rows and columns represent genes and individual cells, respectively. Relative expression levels are shown in yellow (high) and magenta (low) colors.

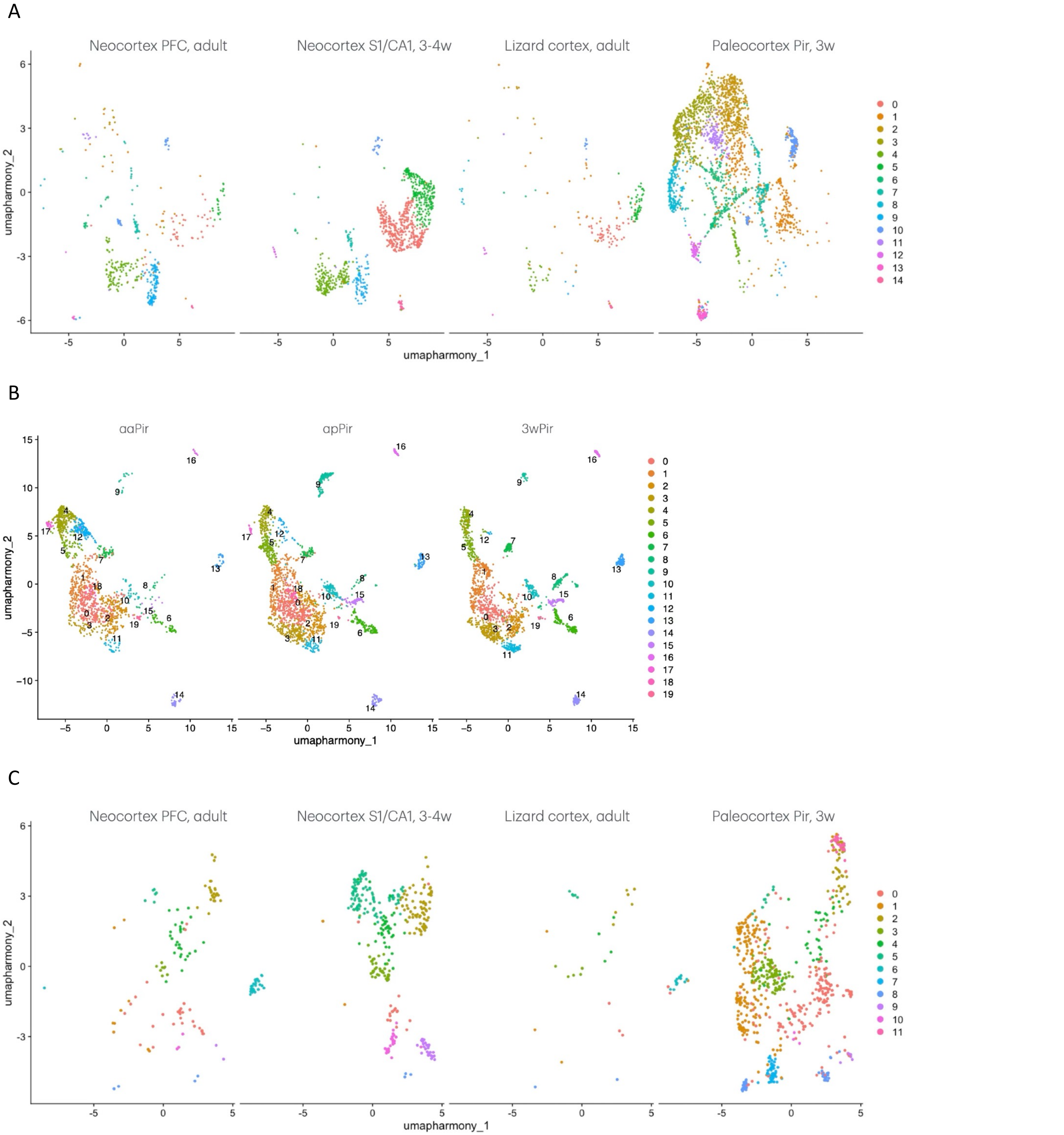


Supplementary Figure 4. Batch-segregated UMAP projections after harmony-based batch correction.
(A) UMAP projection of glutamatergic clusters (Cq0-14) across the four inter-cortical and interspecies datasets. (B) UMAP projection of paleocortical glutamatergic clusters (Cp0-19) across three piriform cortex datasets (aaPir, apPir, and 3wPir). (C) UMAP projection of GABAergic clusters across the four inter-cortical and interspecies datasets (Cg0-11). Clusters were color-coded according to the Seurat identity.

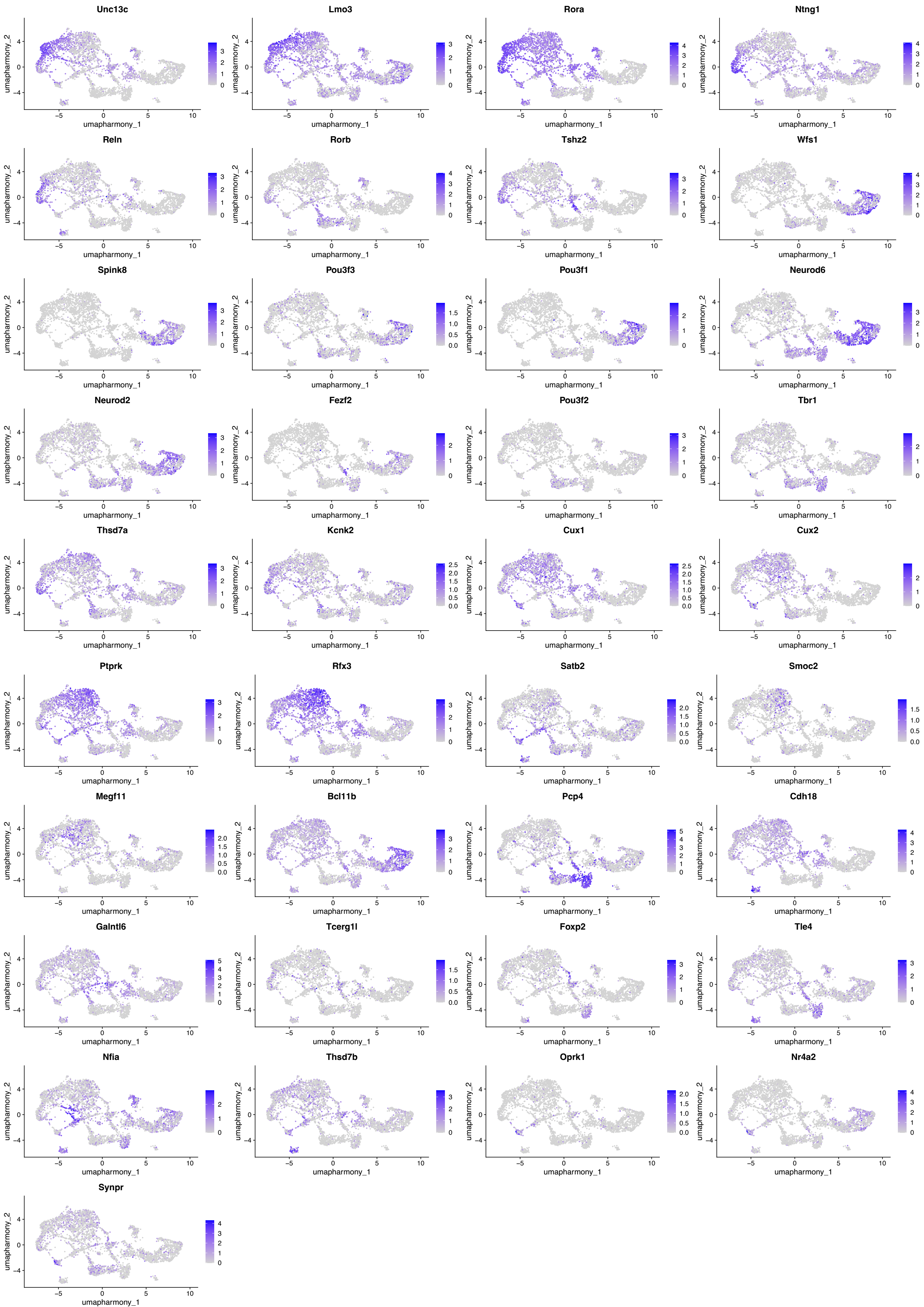
Unc13c
Lmo3
Rora
Ntng1
Reln
Rorb
Tshz2
Wfs1
Spink8
Pou3f3
Pou3f1
Neurod6
Neurod2
Fezf2
Pou3f2
Tbr1
Thsd7a
Kcnk2
Cux1
Cux2
Ptprk
Rfx3
Satb2
Smoc2
Megf11
Bcl11b
Pcp4
Cdh18
Galntl6
Tcerg1l
Foxp2
Tle4
Nfia
Thsd7b
Oprk1
Nr4a2
Synpr
umapharmony_1
umapharmony_2

Supplementary Figure 5. Gene expression patterns of representative DEGs in the the integrated datasets. Dots indicate glutamatergic neurons expressing the indicated genes in the UMAP space across the four inter-cortical and interspecies datasets. Relative expression levels are shown in yellow (high) and magenta (low) colors.

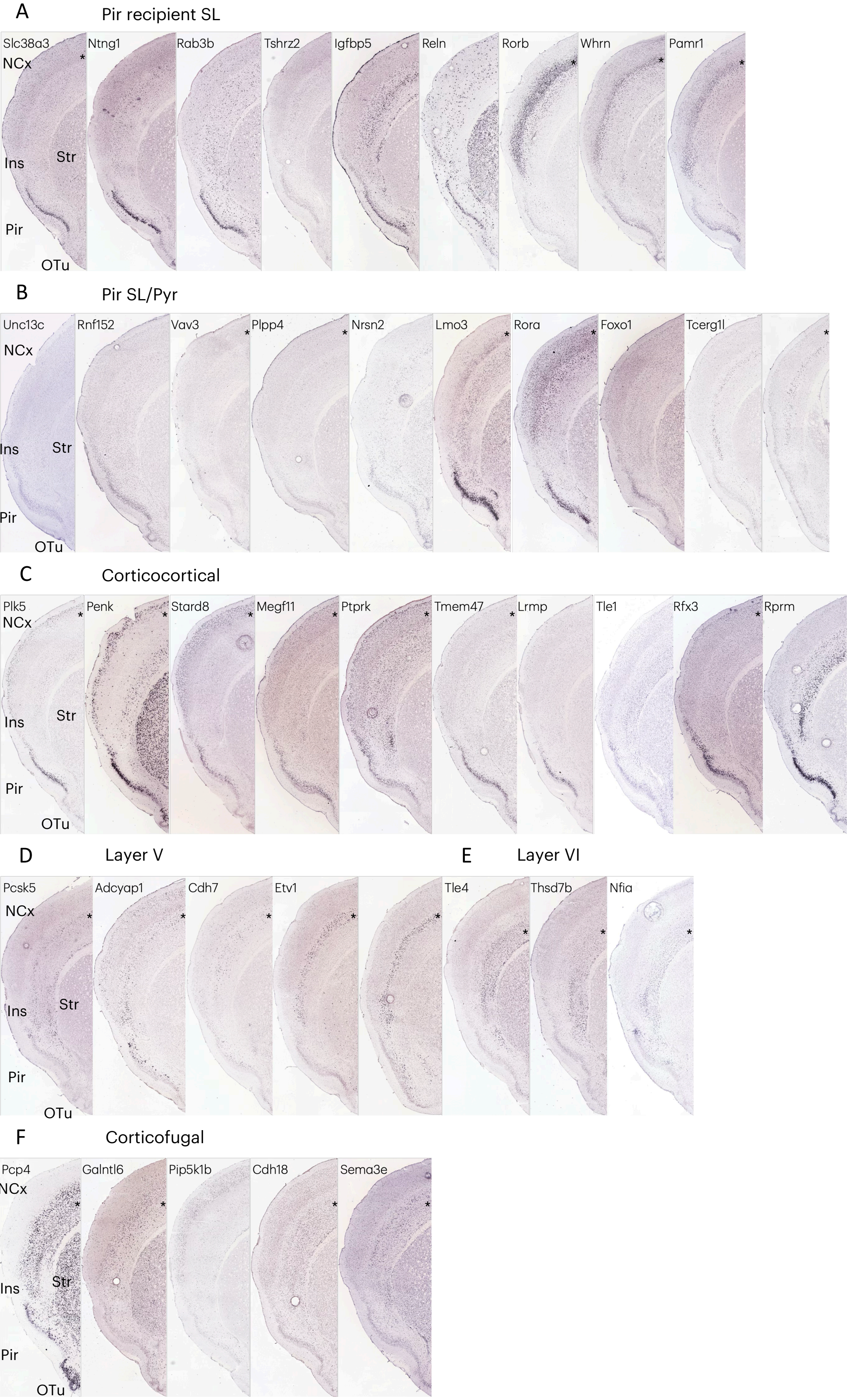
A
Pir recipient SL
Slc38a3
Ntng1
Rab3b
Tshrz2
Igfbp5
Reln
Rorb
Whrn
Pamr1
NCx
Ins
Str
Pir
OTu
B
Pir SL/Pyr
Unc13c
Rnf152
Vav3
Plpp4
Nrsn2
Lmo3
Rora
Foxo1
Tcerg1l
NCx
Ins
Str
Pir
OTu
C
Corticocortical
Plk5
Penk
Stard8
Megf11
Ptprk
Tmem47
Lrmp
Tle1
Rfx3
Rprm
NCx
Ins
Str
Pir
OTu
D
Layer V
Pcsk5
Adcyap1
Cdh7
Etv1
NCx
Ins
Str
Pir
OTu
E
Layer VI
Tle4
Thsd7b
Nfia
F
Corticofugal
Pcp4
Galntl6
Pip5k1b
Cdh18
Sema3e
NCx
Ins
Str
Pir
OTu

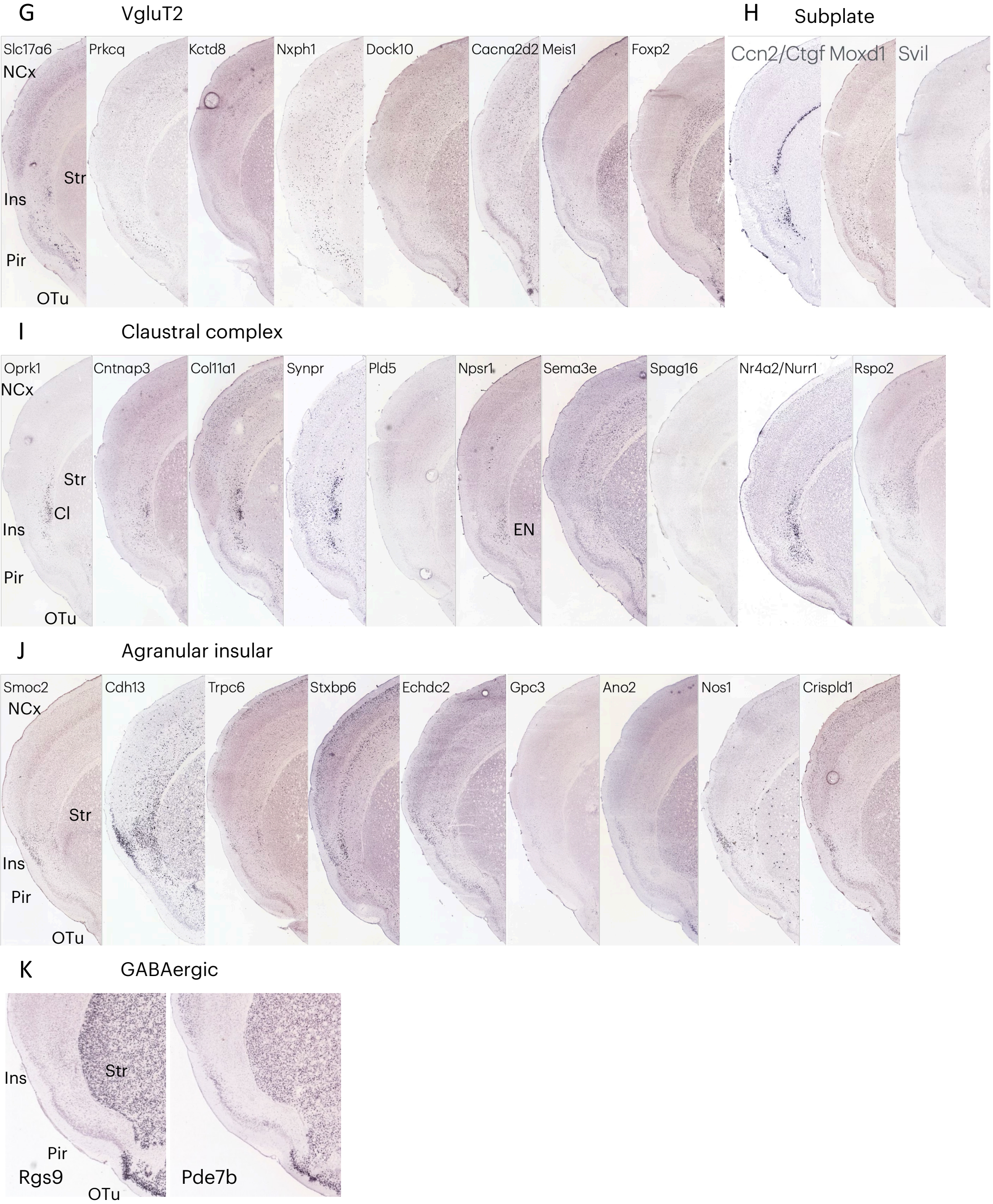


Supplementary Figure 6. Spatial expression patterns of representative DEGs across cortical and subcortical regions based on the Allen Brain Atlas.

(A–F) Layer-specific expression patterns across piriform layers II and III. Asterisks indicate the corresponding expression in the neocortex.

(G) Expression patterns of *Vglut2*-associated genes.

(H–K) Expression in the subplate, claustral complex, insular cortex, and paleocortical GABAergic neurons. Abbreviations are as in the main text.

(A) Genes expressed in the layer IIa. (B) Genes expressed in layers IIa and IIb. (C) Genes expressed in the layer IIb. (D) Genes expressed in the layer IIIa. (E) Genes expressed in the layer IIIb. (F) Genes expressed in layers IIIa and IIIb. (G) *VgluT2*-related genes. (H) Genes expressed in the subplate. (I) Genes expressed in the claustral complex. (J) Genes expressed in the insular cortex. (K) Genes expressed in the paleocortical GABAergic neurons. Abbreviations: Cl, claustrum; EN, endopiriform nucleus; Ins, insular; Pyr, superficial pyramidal; SL, semilunar; SP, subplate.

### Supplementary Table 1. DEG-based correspondence between Cqs and Cps.

| Subtype | Cp | Cq | Marker |
|---|---|---|---|
| Pir-dominant neuron markers | 1,4,5,12,17 | 3,8 | Unc13c (Cq3/8 and Cp1/5), Ntng1 (Cq8 and Cp4/5/12/17), Grik1 (Cq1/8 and Cp4/5) |
| Corticocortical neuron markers | 0,2,3,11 | 2,11 | Megf11 (Cq11 and Cp0), Smoc2 (Cq2/12 and Cp11), Rfx3 (Cq2/11 and Cp0/2/3/11) |
| Corticofugal neuron markers | 8,10,15,16,19 | 1,6,7,13 | Nxph1 (Cq1/7 and Cp8/16), Galntl6 (Cq1/7 and Cp8/15), Pcsk5 (Cq6/13 and Cp6/10),<br>Nfia (Cq6 and Cp6/10/19), Zfhx3 (Cq7 and Cp8/16), Cacna2d2 (Cq7 and Cp8), Tcerg1l (Cq7 and Cp5/8) |
| Claustral neuron markers | 14 | 12 | Oprk1 (Cq12 and Cp14/16), Nr4a2 (Cq12 and Cp6,13,14), Synpr (Cq12 and Cp14) |
